# Identification of selective sweeps in bacteria

**DOI:** 10.1101/2023.03.16.533065

**Authors:** Oren Avram, Eli Levy Karin, Jukka Corander, Yaara Oren, Tal Pupko

## Abstract

Selective sweeps occur when a beneficial mutation spreads rapidly throughout the population due to natural selection. Searching for selective sweeps has proved to be one of the most fruitful ways to detect the footprints selection leaves on the genome. With a plethora of detection tools, the study of selective sweeps in eukaryotic systems is a well-established field of research. However, the search for fragment-specific selective sweeps among bacterial strains received little to no attention so far. In our work, we demonstrate that inter-strains locus-specific selective sweeps can be detected in bacteria. We introduce the *SINCOPA* algorithm, the first phylogeny-based method for soft and incomplete selective sweeps detection. We use *SINCOPA* to explore inter-strains locus-specific selective sweeps in a dataset containing more than 500 microbial genomes. We observe strong evidence in several loci for locus-specific selective sweeps including genes involved in biofilm formation and others that are related to coping with various unfavorable environmental conditions. *SINCOPA* is freely accessible as a user-friendly web server application at https://sincopa.tau.ac.il/.

## Introduction

Since the first genome of *Haemophilus influenzae* Rd was sequenced^1^, bacteriology has changed dramatically. During the two and a half decades that have passed, sequencing technologies have been improved, and sequencing prices have become several folds cheaper. Nowadays, there are thousands of publicly available full-genome sequences of many bacterial species^2^. In addition, it is currently routine to sample and sequence numerous individual genomes of any single bacterial species under study, paving the way for novel bacterial population genetics studies^3,4^. One important goal in population genetics is to detect the signature of adaptive selection. In eukaryotes, given genomic data, a common way to detect adaptations is to look for footprints of selective sweeps that are driven by positive selection^5^. Selective sweeps occur when a beneficial mutation spreads rapidly throughout the population due to natural selection^5^. The hallmark of selective sweeps is reduced genetic variation in the locus of the mutation and its flanking regions (see illustration in Supplementary Figure S1). Searching for selective sweeps has proved to be one of the most fruitful ways to detect the footprints selection leaves on the genome. There is a plethora of sophisticated detection methods for eukaryotic data analysis^6–9^. Additionally, there is a variety of eukaryotic genes that were shown to undergo selective sweeps, from drug resistance in Protista^10^ to brain size regulation in Mammalia^11^.

Although the study of selective sweeps in eukaryotic systems is a well-established field of research^12^, they have not been studied in non-eukaryotes, with the exception of few notable examples^13–18^. One possible reason for the lack of studies aiming to detect selective sweeps in bacteria is the common working hypothesis that once a bacterium gains an adaptive mutation, clonal expansion is likely to happen, i.e., the allele with the entire genome will be fixed or nearly fixed in the population (sometimes referred also as genome-wide selective sweeps^13,14^). However, in this study, we postulate that an increase in frequency by recombination may also occur even across different species if the selectively advantageous allele is repeatedly integrated into the genomes of members of the bacterial community, via lateral fragment transfer (LFT). The term fragment, rather than gene, may reflect either coding or non-coding DNA (e.g., regulatory region). Thus, if LFT is common enough, a DNA fragment can increase in frequency without a significant increase in the frequency of the harboring genome, i.e., a bacterial fragment-specific selective sweep. This concept challenges the paradigm in which the increase in allele frequency is driven by clonal expansion. We test for the existence of selective sweeps in 541 bacterial genomes spanning four different species. To this end, we developed the first phylogeny-based algorithm for selective sweeps detection tailored to bacterial data. Next, we applied our algorithm to a large dataset of dozens of *Enterobacteriaceae* genomes from multiple lineages. Our results clearly showed evidence of fragment-specific selective sweeps, e.g., in genes involved in ions detoxification and biofilm formation. Then, we extended our analysis to three other datasets, each containing more than a hundred bacterial genomes, with species that exhibit varying recombination rates. Our results suggested that bacterial adaption can occur quickly by recurrent integration of beneficial alleles mediated by inter-strain LFT.

## Results

We have previously studied lateral transfer of regulatory regions within bacteria^19^. In that study, we curated a dataset of multiple sequence alignments (MSAs) composed of orthologous genes from various *Enterobacteriaceae* species. Upon detailed inspection, a few MSAs puzzled us greatly, for example, the MSA of the *hemH* locus. Shown in Figure 1 is a section from the MSA of the regulatory region of the *hemH* gene. This MSA is roughly 200 bps long and encompasses 72 strains. It is clear that two different allele types exist in this locus: an allele with a gap and an allele without one (herein referred to as the “gap allele” and the “non-gap allele”, respectively). We mapped the strains harboring the two allele types onto the species phylogeny and observed that the spread of the two alleles is highly incongruent with the species phylogeny (neither the group of strains harboring the gap allele nor the group harboring the non-gap allele was monophyletic), a pattern suggesting lateral transfer. We reconstructed the local maximum likelihood (ML) tree using RAxML^20^ by providing this MSA section. We then compared it to the species tree, which was reconstructed based on the species’ core proteome; see Methods. Based on the AU test^21^ as implemented in CONSEL^22^, we could reject the species tree from the confidence set of trees for the relevant MSA segment (P < 1e-35). This suggests the incongruence between the local tree and the species topology is statistically significant. There are three possible explanations for such a pattern. First, there may have been three independent deletions in the lineages leading to strains harboring the gap allele. This is highly unlikely as the gaps in all gap-allele orthologs are in the exact same position. The second possibility, which is also unlikely, is the occurrence of three independent *de novo* fragment generations in the non-gap allele. The third and most likely explanation points to several LFT events. Upon closer examination, we noticed a puzzling pattern of conservation of the non-gap allele. Figure 2 shows the same MSA with rearranged rows, where all the non-gap orthologs are relocated to the upper region. While the gap allele is genetically variable, the non-gap allele has almost no variation: average nucleotide pairwise distance of 0.0005 and 0.0205 substitutions per site, for the non-gap allele and the gap allele, respectively (P < 0.0001; t-test).

**Figure 1.**
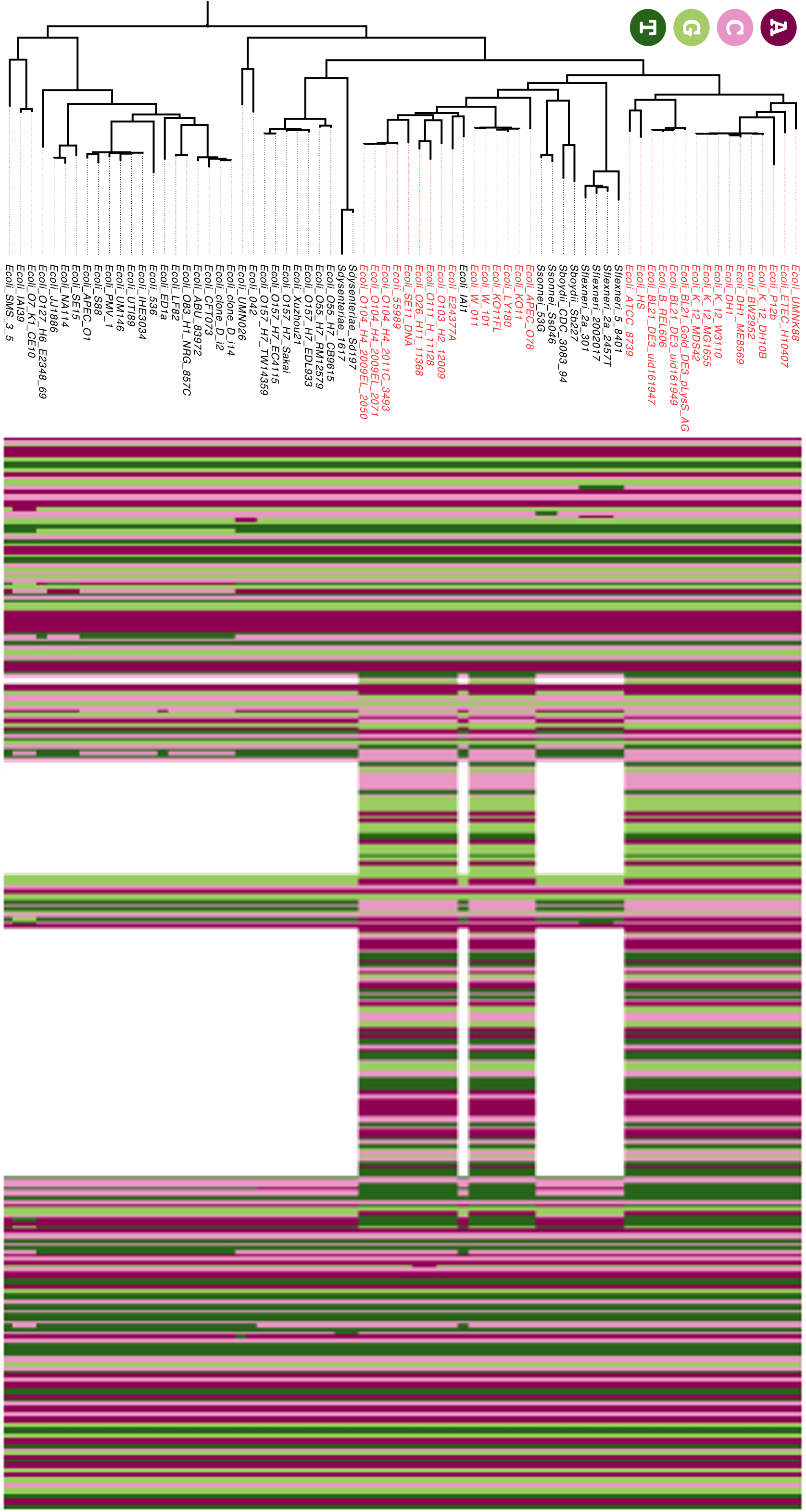
A segment from the MSA of the regulatory region of the *hemH* gene in *Enterobacteriaceae*. This segment is roughly 300 bps wide and it encompasses 72 *Enterobacteriaceae* strains. Each of the four colors represents a different nucleotide and gaps are represented in white. Each line corresponds to the strain from its left. There are two allele types for this locus: an allele with a gap and an allele without one. The spread of these two alleles is incongruent with the species phylogeny (the strains with the non-gap allele are colored in red). The phylogenetic pattern together with the sequence similarity of the red species group suggest a partial sweep event within the non-gap allele group (see also Figure 2).

**Figure 2.**
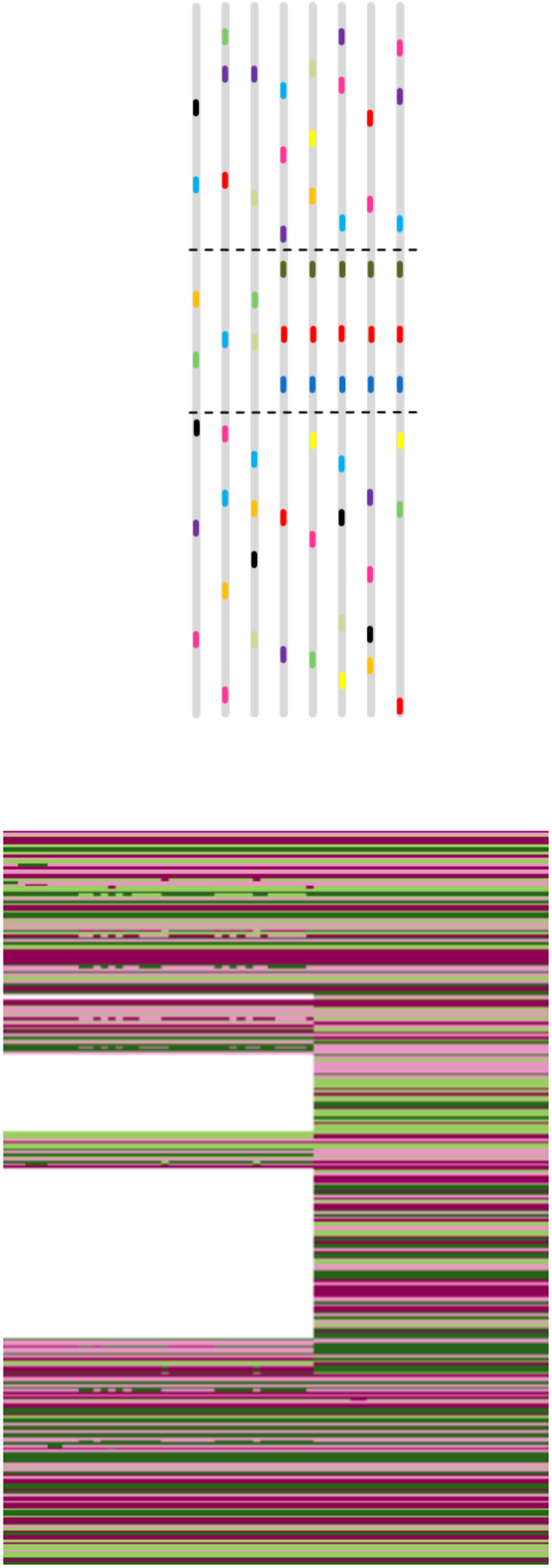
A (rearranged) partial selective sweep illustration (Panel A) and a (rearranged) putative partial selective sweep in empirical data (Panel B). Panel A shows an illustrated example of homologous loci after a selective sweep event. Panel B shows the same MSA from Figure 1 after rows rearrangement (all the gap alleles are clustered at the bottom). Notice the similarity between the illustrated swept region (within the two dashed lines in Panel A) and the empirically aligned region of the *Enterobacteriaceae* locus (Panel B). The underlying hypothesis is that the adaptive allele originated in one of the species in the upper group (see purple strains in Figure 1) and spread among these strains eliminating any genetic variance encountered.

What could explain this 41-fold difference in the accumulation of mutations between the two alleles? We postulated that the observed sequence pattern may have resulted from a partial selective sweep rendered by LFT. Upon careful examination of the two allele groups, the conservation inconsistency across the suspected region, together with the convergence of the sequence variability levels in the flanking area of the suspected region, implies that the LFT occurred multiple times independently and at a high rate, a pattern that is consistent with a selective benefit for that allele (Figure 2).

We next aimed at identifying additional loci in the *Enterobacteriaceae* dataset that show similar patterns congruent with selective sweep events. To this end, we examined several previously developed selective sweep detection methodologies such as SWEEPFINDER^7^, *iHS*^8^, and *nS*_*L*9_. These methods, however, were designed to detect selective sweeps in eukaryotes and were shown to be both sensitive and accurate when applied to eukaryotic datasets. However, these methods cannot be directly applied, if at all, to bacterial data such as the ones analyzed in this study for several reasons. First, they do not handle insertion-deletion (indel) events, and this is reasonable as indels are not common within a eukaryotic species as their genomes are less prone to perturbations relative to bacteria^23^. Second, these methods assume polarity, that is, knowledge regarding which allele is ancestral, and which is derived. Polarity inference is straightforward in eukaryotes (based on an outgroup sequence) since horizontal transfers are either absent or extremely rare. However, in bacteria, this is clearly not the case. Third, these methods do not take the phylogeny into account^24^, and thus, may give spurious selective sweep detection, for example, upon examining an MSA containing two monophyletic groups that underwent ancient speciation.

Considering these gaps, we were motivated to develop an algorithm for inferring fragment-specific selective sweeps tailored to bacterial data. The following features were taken into account in the design: (1) The algorithm should consider the extent of expected sequence divergence observed among different bacterial strains (to avoid spurious detection of sweep events) and should search for discrepancies in the sequence patterns from those expected based on the phylogenetic species tree; (2) The algorithm should detect conservation pattern within the suspected horizontally transferred region, similar to methods designed to analyze eukaryotic data^7^, i.e., a region with a single highly conserved allele (in case of partial sweeps or more than one conserved allele in case of soft sweeps) in some clades of the tree, suggesting a putative rapid spread of the beneficial allele; (3) The algorithm should provide a single score, reflecting the propensity that a sub-region within a given MSA experienced a selective sweep. (4) The algorithm must be extremely fast to allow analyzing (at least) dozens of fully-sequenced bacterial genomes.

To this end, we devised the *Similarity of INCOngruent PAtterns* (*SINCOPA*) algorithm – the first phylogeny-based algorithm for selective sweeps detection in bacteria. *SINCOPA* algorithm is deployed on a user-friendly web server and is freely available at https://sincopa.tau.ac.il. The input of *SINCOPA* is an MSA and a species phylogeny. The output of the algorithm is a sweeps scores vector, where each score corresponds to a specific window (i.e., a set of consecutive columns) in the alignment. The algorithm is divided into two main steps. In the first step, homoplastic columns are identified, that is, columns that are incongruent with the input tree. Specifically, given a set of aligned sequences corresponding to a set of species and the topology of their species tree, the cost *c* for each alignment column, i.e., the minimum number of evolutionary changes required to explain the data, is computed using the classic Fitch dynamic-programming algorithm^25^. Then, the column is marked as homoplastic only if *c* + 1 is larger than the number of different DNA characters occurring in the current column. This definition of homoplastic positions is the same as in other phylogenetic studies^26^. Gaps are taken into account as a (fifth) character. This computation is a first step towards a rapid identification of regions within the alignment that are highly likely to have experienced horizontal transfer. However, not all horizontal transfers are associated with selective sweeps. It may be that a horizontal transfer event occurred neutrally, and in which case we expect that the spread of the transferred allele was relatively slow, and it would be accompanied by the accumulation of mutations within the transferred allele. In contrast, a selective sweep would result in little variation (if at all) within the transferred allele. In the second step of the algorithm described below, we compute a sweep statistic, 0 ≤ *S* ≤ 1, which is tailored to distinguish between these two cases. Large values of *S* indicate similarity between the allelic patterns of different column positions, suggesting that horizontal transfer was rapid relative to the accumulation of mutations, whereas low *S* values indicate cases where there was no horizontal transfer or horizontal transfer that was neutral or nearly neutral.

To search for regions that are both enriched with homoplastic positions and that the homoplastic positions share similar patterns with each other, we use a sliding window of length *w* (50 by default) with a stride of one. The *w* positions in each window include both homoplastic and non-homoplastic positions. In each window, all pairs of homoplastic columns are enumerated and an association score, *α*, for each such pair of homoplastic columns is calculated (*α=0* for all other pairs, namely, pairs of at least one non-homoplastic column). A high value of *α* suggests that the two columns are of a similar pattern. A greater number of high-*α*-values within a window indicates a higher chance that the entire region experienced a selective sweep. Formally, the *S* statistic of a window is defined as follows:

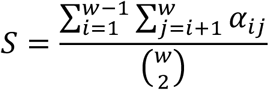

where *α*_*ij*_ is the *α* score between columns *i* and *j* within the window. For the sake of efficiency, only the first window is fully computed. To compute any other window, the contributions of the column pairs involving the two non-overlapping columns (i.e., the last column of the window in question and the first column of its previous window) are being computed (and added or removed, respectively). This way the running time complexity is *O*(*w*^2^ + *nw*), where *n* represents the length of the MSA.

The procedure for calculating the association score *α* between two alignment columns we suggest is straightforward. Intuitively, for each nucleotide in the first (reference) column we search for the most frequent matching nucleotide in the second column. The highest score is obtained when each nucleotide in the first column is matched to exactly one nucleotide in the second column. A low score is obtained when the association between the nucleotides in the two positions is low. In order to keep the score symmetric, we repeat the above computation twice (once for each column as a reference) and take the average. Finally, we get *α* by normalizing the averaged result by the column length (thus, the score of different alignments with a different number of species is comparable). A pseudo-code that describes in detail this computation is provided together with a detailed computation on a toy example (Supplementary Figure S2).

### Searching for sweeps in the *Enterobacteriaceae* dataset

We used as input the inferred *Enterobacteriaceae* ortholog groups and the corresponding core-proteome-based species tree provided by M1CR0B1AL1Z3R^27^ (*E. fergusonii* ATCC 35469 was used to root the tree). We extracted each ortholog group with its corresponding promoter and aligned it using MAFFT^28^. For each MSA, the *SINCOPA* algorithm expects a corresponding species phylogeny. In the case of a core gene ortholog group, we used the same (core-proteome-based) species tree. For a non-core gene ortholog group (i.e., an ortholog group in which some species lack a ‘member gene’), we adjusted the species tree by trimming it to contain only the relevant species of the MSA in question. Next, each MSA and its corresponding species tree were used as input to *SINCOPA*. In order to examine how prevalent sweeps are in the whole dataset, we defined an MSA-level statistic *S*\*, as the highest *S* statistic obtained across all of its windows. We then examined the distribution of the *S*\* statistic across all ortholog groups. We defined candidate genes that may have undergone selective sweeps using an “outlier approach”, similar to other (eukaryotic) selective sweeps detection methods^8,9^. That is, we were focusing on the top percentile of the abovementioned distribution. The top-hit genes and the *S*\* distribution of all MSAs are shown in Table 1 and Figure 3, respectively.

**Table 1.**
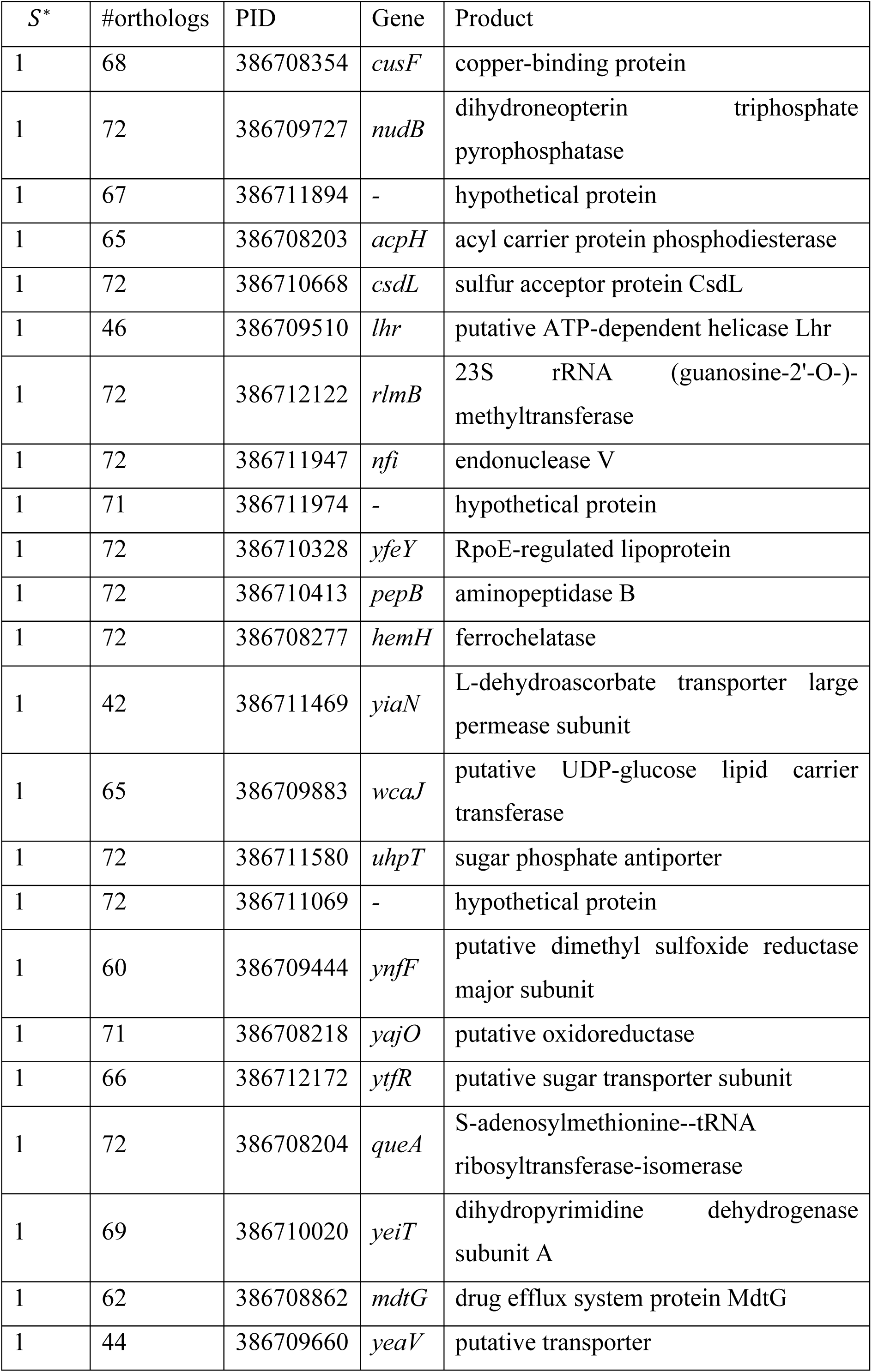
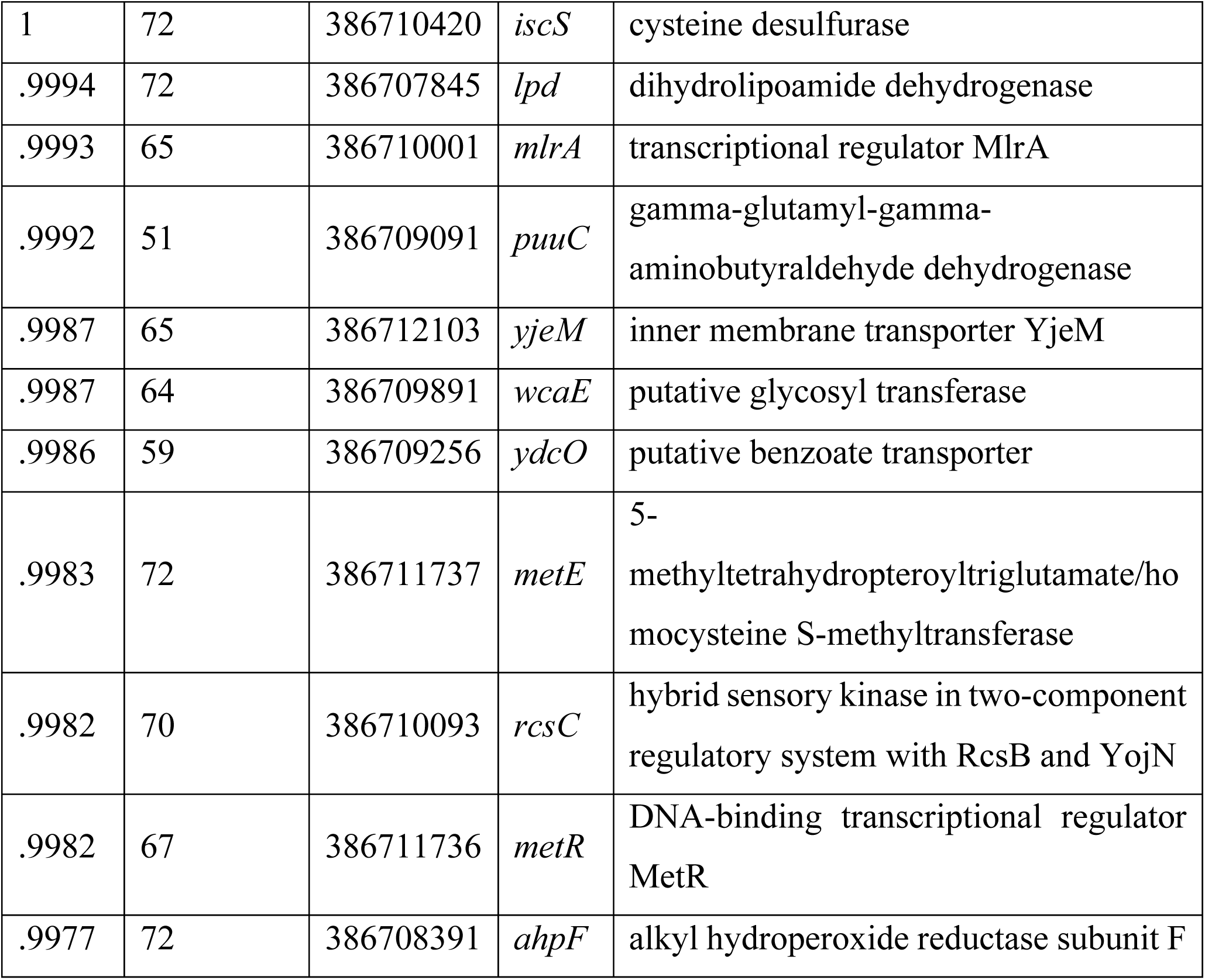
Top *S*\* scoring genes obtained in the *Enterobacteriaceae* dataset analysis.

**Figure 3.**
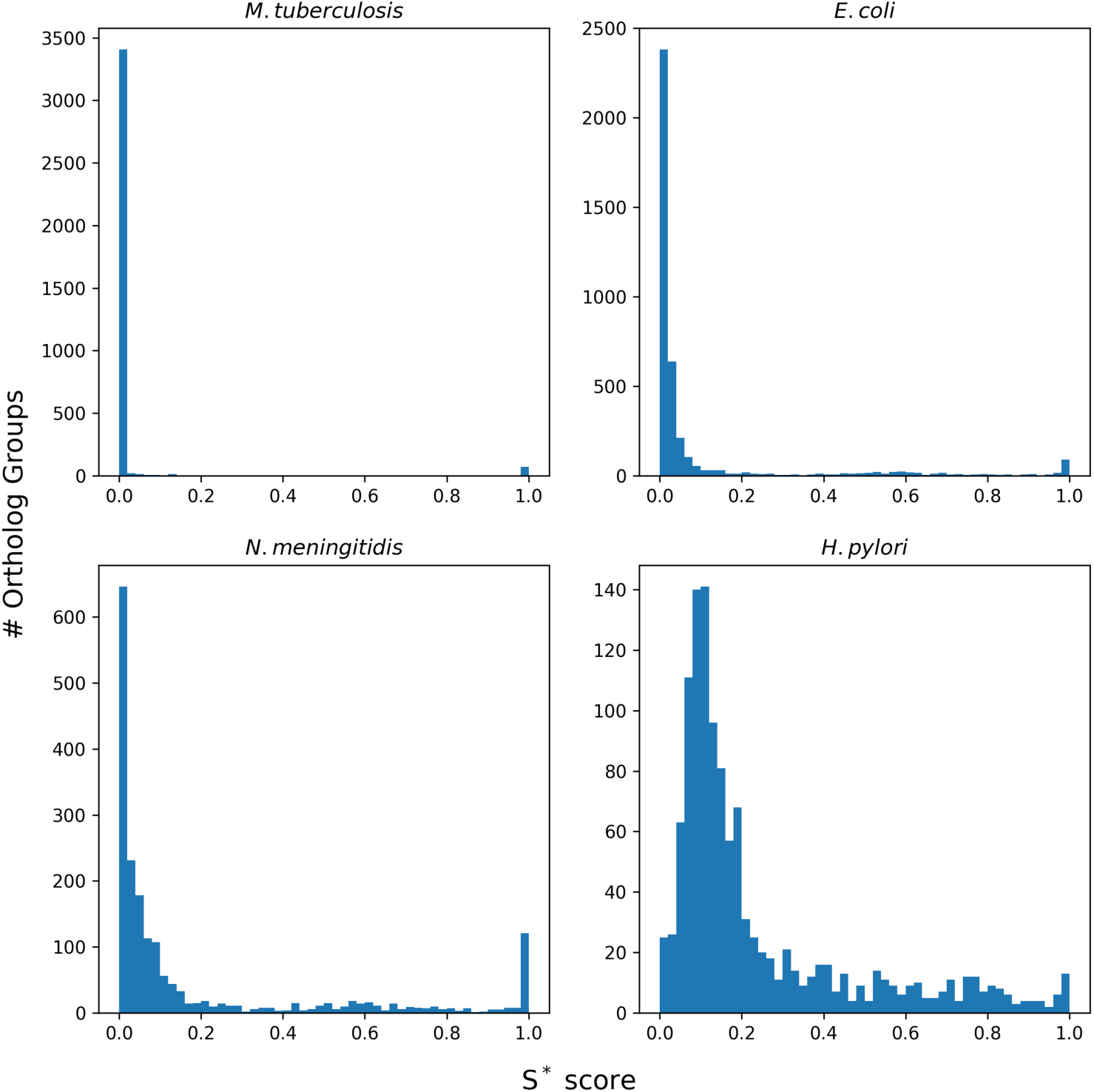
*S*\* scores distributions of the four different datasets analyzed in this study. Shown is the *S*\* distributions of *M. tubercolosis, Enterobacteriaceae, N. meningitidis*, and *H. pylori*. The recombination rates of these four species ascend from *M. tuberculosis* with the lower rate to *H. pylori* with the highest rate. The higher the recombination rate the higher the sweeps ‘signal’ (featured by right-skewness).

Among the top results in *E. coli* we could find the *hemH* gene. This gene was previously shown to undergo regulatory region horizontal transfer. There are two variants of the promoter regions of this gene, each harboring a different transcription factor binding site^19^ (Figures 1 and 2). In this case, it is suggested that the regulatory region horizontal transfer allows rapid switching between two regulatory modes, and thus selection played a major role in the evolution of this region. The low genetic variation in only one of the alleles (the non-gapped allele) suggests that the swept allele spread rapidly among the different strains. Another top-scoring result was the *cusF* gene that is involved in the detoxification of copper ions in *E. coli*^29^. An adaptive allele may confer an advantage in an environment that is contaminated with this metal. Another top hit was *wcaJ*. This gene plays a major role in the assembly of colanic acid exopolysaccharide^30^, which in turn, provides a protective barrier against unfavorable conditions, such as an acidic environment^31^ and extreme drought and high temperatures^32^.

### Extending the sweeps analysis to other bacterial datasets

We further extended our analysis and added another 469 genomic sequences of three other species of different lineages: *Mycobacterium tuberculosis* (182 genomes), *Neisseria meningitidis* (108 genomes), and *Helicobacter pylori* (179 genomes). These bacteria vary in their recombination to mutation ratios^19^. Here again, we established a database for each of these species similarly to the *Enterobacteriaceae* dataset (see Methods). Briefly, we reconstructed each species phylogeny and inferred an orthologs table. Next, we applied *SINCOPA* over each ortholog group in these datasets and examined the *S*\* distribution of each individual species. Figure 3 demonstrates the *S*\* of the datasets of the four species analyzed in this study. As we expected, we could detect a positive correlation between the recombination rates and how skewed the *S*\* distribution is, as detected by *SINCOPA*. While *M. tuberculosis* shows typically zero recombination rate, its *S*\* distribution is only slightly skewed to the right. *E. coli* has a low-moderate recombination rate and indeed shows more right skewness. Accordingly, *N. meningitidis* recombination rate is moderate-high and its *S*\* distribution is further right-skewed. Lastly, the distribution of *H. pylori* is the most right-skewed distribution among the analyzed datasets with agrees with its high recombination rate.

## DISCUSSION

In this work we devise the first phylogeny-based algorithm for detecting fragment-specific selective sweeps in bacterial genomics data. The idea for the study arose after we detected, by visual inspection, several interesting examples of putative sweep patterns in microbial MSAs. This prompted us to conduct a large-scale study to see how prevalent these patterns are and how recombination rate might affect them. For that we developed the *SINCOPA* algorithm and conducted a large-scale study spanning over 500 bacterial genomes with varying recombination rates. We started by investigating an *E. coli* dataset containing 72 genomic sequences. One of the challenges in this work was to establish reliable ortholog groups across dozens of bacterial genomes, identify core genes, and reconstruct a species tree. To this end, we have developed M1CR0B1AL1Z3R^27^ that automates these tasks. The resulting output was used as input to *SINCOPA*, which searches for selective sweeps. We were able to detect a set of genes in which the sweeps signal is at its extremity. Among them were genes that are related to copper ion detoxification and adverse environmental conditions, such as acidity, drought, and high temperature. Next, we extended our study to another 469 genomes of three other bacterial strains with different recombination rates. We detected a positive trend between the sweeps signal detected and the recombination rate documented in the literature for the examined strains. Since this method is not strain-specific but rather general, it can be applied to many other bacterial species. An interesting direction for future research will be to study a dataset of bacterial populations of the same species that (preferably, recently diverged and) inhabit different niches. This way, inferring a selective sweep can be interpreted as an ecological differentiation.

The method we introduced has, however, several limitations. First, due to the logic underlying *SINCOPA*, it cannot detect hard selective sweeps. In contrast to soft/partial selective sweeps, hard sweeps are defined as sweep events in which the sweeping allele reaches fixation. *SINCOPA* seeks homoplastic MSA columns, i.e., positions that show incongruency with the species phylogeny, which are, by definition, non-homogenous. Hence, it will not discriminate between homogenous columns (as they are always consistent with the species tree) that are (1) a consequence of evolutionary-conserved regions and (2) a consequence of a recent hard selective sweeps event. Therefore, this issue, unfortunately, is not addressable via *SINCOPA*. A second limitation is the lack of a coherent simulation platform. Since there is no appropriate statistical model underlying the sweeps hypothesis, it is unclear how to determine the level of false positive predictions (a region that was falsely classified as if it experienced a sweep event) and false negative predictions (a region that was swept but was missed in the analysis). Thus, an ‘outlier approach’ was taken here. We could not find a suitable tool for simulating the evolutionary dynamics underlying multiple bacterial strains from the same species. We did try to back up our conclusions by devising a simulation framework that aims to reflect microbial genes evolution, accounting for horizontal transfers (recombination), mutations, selection, and genetic drift. The simulation pipeline was based on the Wright-Fisher model^33,34^ and extended a previously developed common simulator^35^. For improved performance, the simulator was implemented in C++ and it is available at the project’s online repository: https://github.com/orenavram/SINCOPA/tree/master/LociSimulator. Briefly, in each simulation step (i.e., bacterial generation) each derived bacterium draws a parent bacterium and “inherits” its genome. Depending on the tunable rate parameters, an event such as mutation, recombination (both within and between populations), and selection for adaptive alleles, may occur. At the end of each simulation, a sample is drawn from each lineage. We believe that this process reflects appropriately what we observe in the four different datasets analyzed in this study. When we simulated without selection and ran *SINCOPA*, we observed that the *S*\* distribution was centered near zero and no cases of high *S*\* were found. Although studied extensively, we were unable to reconstruct any (simulation-based) *S*\* distribution that reflects the strong signal we observed in the (empirical) bacterial datasets analyzed in this study (not shown), suggesting that an even more complex model underlies the pattern we observed. An interesting direction for future research will be to devise a probabilistic model of sequence evolution that will generate patterns of *S*\*distributions that reflect those found in empirical datasets.

## Methods

### Establishing a comparative sequence database of bacterial genomes

Our initial analysis included 72 fully sequenced *Enterobacteriaceae* genomes from the Gallery of M1CR0B1AL1Z3R web server^27^. These genomes included 62 *E. coli* genomes and 10 *Shigella* genomes that are obligate intra-intestinal pathogens belonging to the *E. coli* species^36,37^. Genome sizes ranged from 3.976 mega base pairs (*E. coli* K12 MDS42) to 5.697 mega base pairs (*E. coli* O26 H11 11368). The full list is provided in the supplemental information as Table S1. We used M1CR0B1AL1Z3R web server to extract the orthologs table for each dataset^27^. A total of 9,565 ortholog clusters were detected in the dataset. Out of which, 7,224 were classified as accessory genes, that is, genes that were missing in at least one of the analyzed lineages. The remaining 2,341 genes were classified as core genes and their corresponding protein sequences were used for phylogeny reconstruction using RAxML^20^ via M1CR0B1AL1Z3R^27^ (Figure 1). For the sweeps analysis, we considered only clusters that contained more than 20 member genes, resulting in a set of 4,190 ortholog clusters. We then extracted the gene sequences of these 4,190 groups along with their promoter, defined as 300 base pairs upstream to the translation start site. This definition is routinely used in bacterial regulatory studies^38,39^. Each cluster (of promoter and gene) was aligned using MAFFT v7.407^40^. Careful manual quality assurance of the resulting MSAs revealed differences in the start/end positions of several genes. These differences can lead to alignment errors at the alignment edges, which in turn might generate spurious signal for a selective sweep. To avoid such alignment errors, we additionally trimmed every alignment from both 5’ and 3’ edges. In each edge, columns were trimmed toward the MSA’s center, until a column without “-” characters (i.e., gaps) was detected. We discarded 19 MSAs that were found to be too short (<300 bps) after trimming, ending with a total of 4,171 MSAs. Among these MSAs, a total of 1,826 characters were ambiguously sequenced, such as ‘D’ (which stands for either ‘A’, ‘G’ or ‘T’) and ‘N’ (any of the four nucleotides). To overcome this uncertainty another clearance step was performed: each ambiguous character was replaced with the major (non-ambiguous) nucleotide character in its column. On average, there were fewer than two ambiguous characters per alignment such that the expected effect of this step on the analyses is negligible. We repeated this whole same analysis process over three other bacterial datasets, namely, *M. tuberculosis* (182 genomes), *N. meningitidis* (108 genomes), and *H. pylori* (179 genomes). All of these were downloaded on January 1, 2021, from the National Center for Biotechnology Information (NCBI) repository available at https://ftp.ncbi.nlm.nih.gov/genomes/refseq/bacteria/. The in-house scripts Python source code written for all the procedures mentioned above are available at https://github.com/orenavram/SINCOPA.

## Acknowledgments

O.A. was supported in part by a fellowship from the Edmond J. Safra Center for Bioinformatics at Tel Aviv University and by a fellowship from the Dalia and Eli Hurvitz Foundation LTD. This research was supported by the Israel Ministry of Science, Technology and Space, Grant #47133 and the Israel Science Foundation (ISF) grant 2818/21.

## Supplemental Information

**Table S1.** The *E. coli* strains that were used in this study. *E. fergusonii* ATCC 35469 was used as an outgroup for the species phylogeny reconstruction.

|  |  |  |
| --- | --- | --- |
| *E. coli* 042 | *E. coli* JJ1886 | *E. coli* O157 H7 TW14359 |
| *E. coli* 536 | *E. coli* K 12 DH10B | *E. coli* P12b |
| *E. coli* 55989 | *E. coli* K 12 MDS42 | *E. coli* PMV 1 |
| *E. coli* ABU 83972 | *E. coli* K 12 MG1655 | *E. coli* S88 |
| *E. coli* APEC O1 | *E. coli* K 12 W3110 | *E. coli* SE11 DNA |
| *E. coli* APEC O78 | *E. coli* KO11 | *E. coli* SE15 |
| *E. coli* ATCC 8739 | *E. coli* KO11FL | *E. coli* SMS 3 5 |
| *E. coli* B REL606 | *E. coli* LF82 | *E. coli* UM146 |
| *E. coli* BL21 DE3 uid161947 | *E. coli* LY180 | *E. coli* UMN026 |
| *E. coli* BL21 DE3 uid161949 | *E. coli* NA114 | *E. coli* UMNK88 |
| *E. coli* BL21 Gold DE3 pLysS AG | *E. coli* O7 K1 CE10 | *E. coli* UTI89 |
| *E. coli* BW2952 | *E. coli* O26 H11 11368 | *E. coli* W 011 |
| *E. coli* CFT073 | *E. coli* O55 H7 CB9615 | *E. coli* W 101 |
| *E. coli* clone D i2 | *E. coli* O55 H7 RM12579 | *E. coli* Xuzhou21 |
| *E. coli* clone D i14 | *E. coli* O83 H1 NRG 857C | *S. boydii* CDC 3083 94 |
| *E. coli* DH1 ME8569 | *E. coli* O103 H2 12009 | *S. boydii* Sb227 |
| *E. coli* DH1 | *E. coli* O104 H4 2009EL 2050 | *S. dysenteriae* 1617 |
| *E. coli* E24377A | *E. coli* O104 H4 2009EL 2071 | *S. dysenteriae* Sd197 |
| *E. coli* ED1a | *E. coli* O104 H4 2011C 3493 | *S. flexneri* 2a 301 |
| *E. coli* ETEC H10407 | *E. coli* O111 H 11128 | *S. flexneri* 2a 2457T |
| *E. coli* HS | *E. coli* O127 H6 E2348 69 | *S. flexneri* 5 8401 |
| *E. coli* IAI1 | *E. coli* O157 H7 EC4115 | *S. flexneri* 2002017 |
| *E. coli* IAI39 | *E. coli* O157 H7 EDL933 | *S. sonnei* 53G |
| *E. coli* IHE3034 | *E. coli* O157 H7 Sakai | *S. sonnei* Ss046 |

**Supplementary Figure S1.**
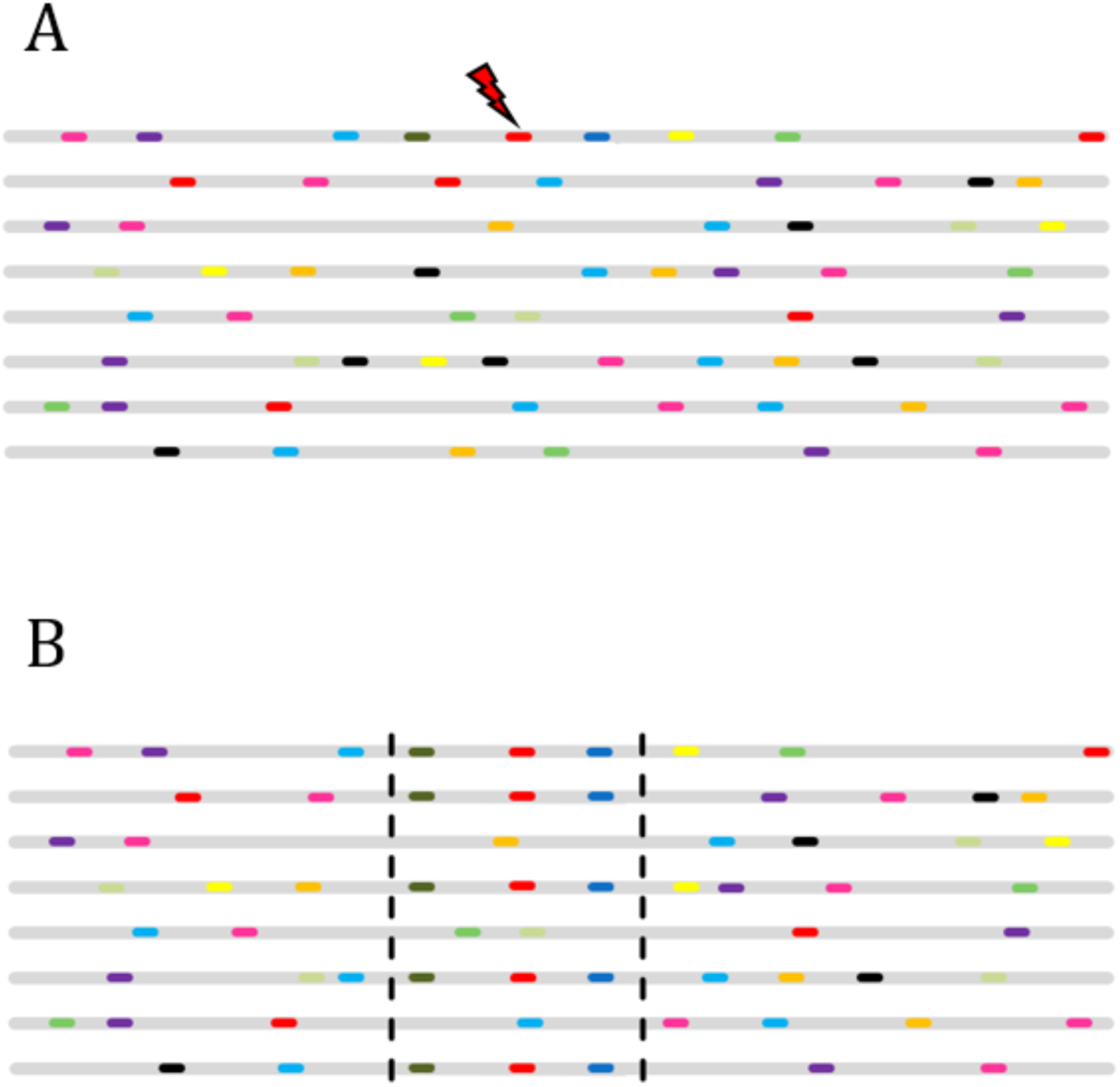
An illustration of the footprint of a selective sweep event on a genomic region. Each of the grey lines illustrates a locus of a single organism and all these loci are homologous to each other. During evolution (Panel A), mutations accumulate (colored dashes). One of the mutations might be beneficial (red lightning) and once it occurs it spreads rapidly (Panel B) throughout the population due to natural selection leaving a footprint of reduced genetic variation (as shown between the two vertical dashed lines). It can be either a de novo mutation or a mutation that was already present in a neutral state, and upon a shift in the environment became adaptive. The presence of increased genetic variation in regions that are more distant from the adaptive mutation is a result of the recombination points between the beneficial allele locus and its flanking regions.

**Supplementary Figure S2.**
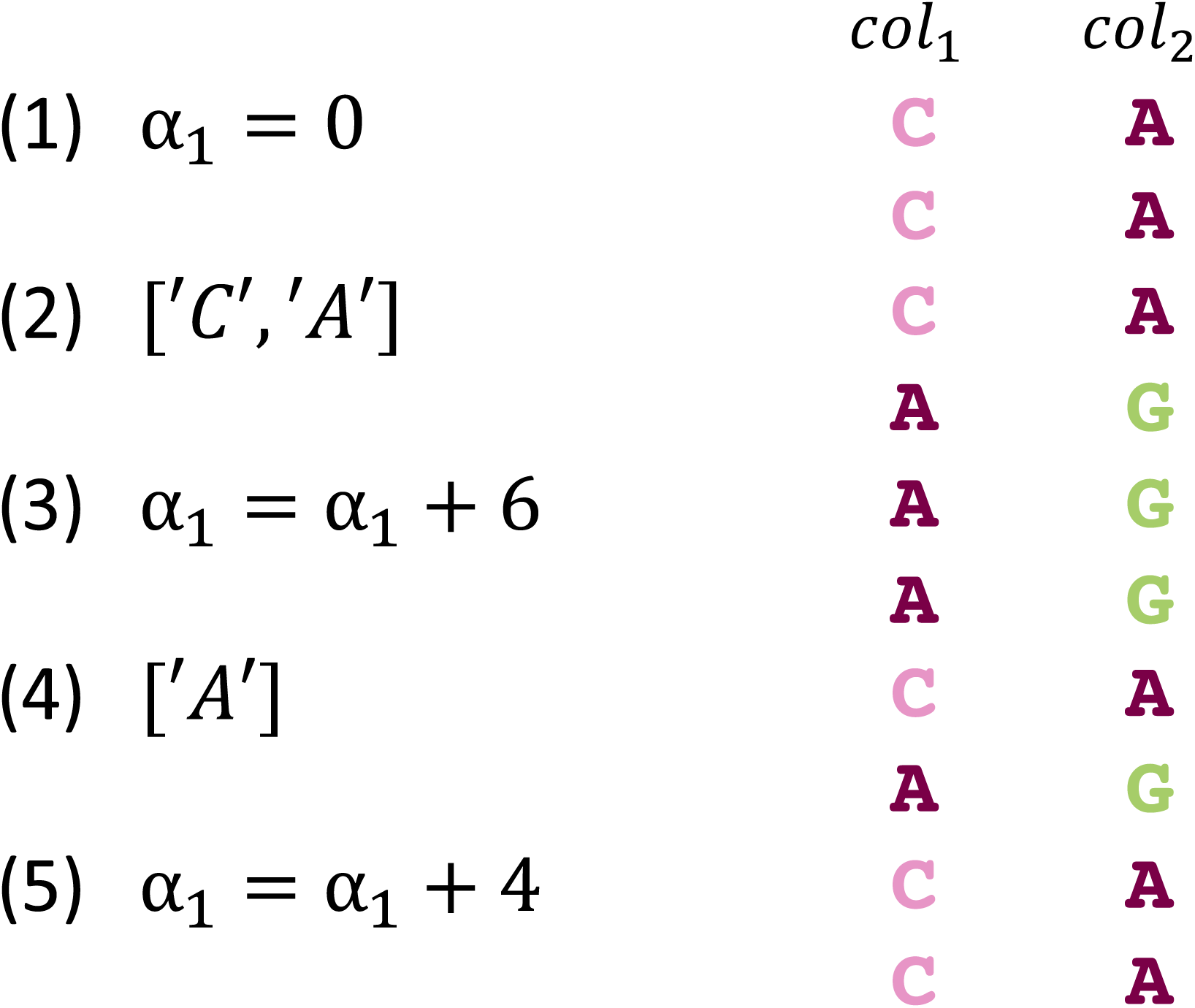
A (partial) illustration of the procedure for calculating the *α* score between two alignment columns. The procedure starts with one alignment column serving as a reference column. First, a counter is initialized to 0 (1). Then, the nucleotides of the reference column (e.g., *col*_*1*_) are sorted by descending frequency (2). Next, for each nucleotide in the reference column, we search for the most frequent matching nucleotide in the other column and increase *α* by the number of matches (3). This process is repeated (4-5) until all the reference column nucleotides were addressed. This procedure is repeated twice (once for each column as a reference column). Let *α*_*1*_ and *α*_(_ represent the scores obtained where *col*_*1*_ and *col*_(_ served as reference columns (respectively), the association score is then given by taking their average.

